# The isoflavonoid brazilin inhibits viability and cell migration in breast cancer cells

**DOI:** 10.1101/2023.08.17.553723

**Authors:** Alberto Hernández-Moreno, Dania A. Nava-Tapia, Jorge Bello-Martínez, Monserrat Olea-Flores, Tadeo Hernández-Moreno, Miriam D. Zuñiga-Eulogio, Napoleón Navarro-Tito

## Abstract

Breast cancer is the most common neoplasm diagnosed in women and is the leading cause of cancer death worldwide. In recent years, compounds isolated from natural sources have been proposed as potential molecules in therapy for breast cancer. In this regard, brazilin has been evaluated in various biological sceneries and has shown pharmacological functions, including anticancer and anti-inflammatory activities. Brazilin was obtained from *Haematoxylum brasiletto*. The chemical structure was confirmed by spectroscopic data (^1^H-NMR, ^13^C-NMR). Concerning biological activity, by MTT assays, brazilin showed cytotoxic effects on MCF7 and MDA-MB-231 breast cancer cell lines. Interestingly, brazilin was not toxic in MCF10A non-tumorigenic breast epithelial cells. We also observed morphological changes to a rounded phenotype associated with apoptosis in breast cancer cell lines and decreased cell migration in a dose and time-dependent manner. By *in silico* analysis, we found that brazilin interacts with JAK1, JAK2, and iNOS, essential molecules driven cell migration and metastasis in cancer. These data suggest that brazilin can potentially be used as an anti-cancer agent in the future.

## 1. Introduction

Breast cancer is the most common neoplasm and the leading cause of cancer death among women aged 45 to 55 years worldwide, with more than two million cases in 2020 [1]. Among all breast cancer types, triple-negative breast cancer (TNBC) is the most aggressive, metastatic, and displays chemoresistance in patients [2]. Some hallmarks of cancer are hyperproliferation, resistance to apoptosis, invasion, angiogenesis, and metastasis [3]. Metastasis is the leading cause of death in cancer patients and is a multi-step process. During metastasis, tumor cells separate from neighboring cells, losing their cell-cell union and cell-extracellular matrix, acquire the migratory capacity, invade the stroma, intravasation to the bloodstream and extravasation in a secondary organ, and grow, forming a secondary tumor [4], [5]. Despite advances in recent years, strategies to counteract cancer progression with the use of mastectomy [6], radiation therapy [7], chemotherapy [8], hormonal therapy, and targeted therapy [9] have not been fully effective since 90% of women who have received these treatments have shown side effects [10,11]. Therefore, new therapeutic agents are needed for the treatment of breast cancer [12], [13].

The diversity of natural products has played and will continue to play an essential role in drug discovery and development [15], [16]. However, they can rarely be used directly in clinical applications due to their low solubility, metabolic instability, and action mechanisms [17]. According to Newman and collaborators, about 75 (41 %) new approved anticancer drugs from 1946 to 2019 were inspired or derived from natural sources [16]. The anticancer effects of numerous flavonoids’ subclasses members have been extensively reviewed, and they can modulate ROS responses, arrest cell cycle, induce cancer cell death, and suppress their proliferation and invasiveness [17] [18]. The Isoflavonoid subclass is a remarkable source of antitumor agents; isoflavonoids such as genistein, daidzein, biochanin A, and glycitein have been taken to the stage of clinical trials for the development of new anti-invasive chemotherapeutics [19].

Brazilin [(6aS, 11bR) -7, 11b-dihydro-6H-indeno [2,1-c] chromene-3, 6a, 9, 10-tetrol], isolated from the ethanolic extracts of the heartwood of *H. brasiletto* (Fabaceae), is a natural tetracyclic homo isoflavonoid product in which a chroman skeleton fused in cis with a 2,3-dihydro-1H indene residue. Previously, brazilin has shown antiproliferative activity in various cancer cell lines such as A549, LS180, H1299, HeLa, SiHa, and MDA-MB-231 [20], [21]. Brazilin also inhibits cell growth and induces apoptosis through caspase-3 activation and subsequent poly(ADP-ribose) polymerase (PARP) cleavage in human U87 glioblastoma cells and U266 myeloma cells [22], and as an inhibitor of P-glycoprotein (Pgp) in MCF7 cells [24]. However, information on the mechanism of action of brazilin on tumor progression is unknown. In the present study, we evaluated the effects of brazilin on cell viability, morphology, and migration in two breast cancer cell lines, MCF7 and MDA-MB-231.

## 2. Materials and Methods

### 2.1 General Information

All materials were analytical grade and used as received unless otherwise noted. Flash chromatography was performed with Silica Flash 60 230-400 mesh. TLC was performed on silica gel 60 F254 pre-coated TLC aluminum sheets (E. Merck, Germany), and two alternative reagents were used for visualization: (a) iodine and (b) KMnO_4_ in water. ^1^H and ^13^C NMR spectra were recorded in CDCl3 on a Bruker AVANCE III HD instrument (500 MHz) using TMS as an internal reference. Chemical shifts (δ) were determined in parts per million, and coupling constant values (J) are given in hertz.

Purification of brazilin: The ethanolic extract from the heartwood powder of *H. brasiletto* was obtained by maceration with EthOH at room temperature for seven days with regular stirring. The combined ethanol extracts were evaporated under reduced pressure on a rotary evaporator to yield the total extract. To the total extract, 250 mL of an aqueous mixture (3: 2 of H_2_O/EthOH) was added, and the resulting suspension was fractionated by liquid-liquid separation with n-hexane, dichloromethane (CH_2_Cl_2_) to produce the corresponding fractions of low and low-media polarity, respectively. Fractions were stored at -4 °C in amber glass vials until use. Brazilin was purified on flash chromatography using gradient elution with CH_2_Cl_2_/EtOAc/MeOH mixtures (9: 1: 0.5), yielding 11 main fractions (3A-3K). Fraction 3B was purified by flash chromatography eluting with n-hexane: EtOAc: MeOH (45: 50: 5) to yield 8 new fractions (4A-4H). Final purification of fraction 4C using EtOAc: CHCl_3_: MeOH (70:30:10 + 50 μL formic acid) resulting in the isolation of brazilin. The ^1^H and ^13^C NMR spectroscopic data are similar to those reported above. (6 a S, 11 b R) -7.11 b -dihydro-6 H -indeno [2,1-c] chromene-3.6 a, 9,10-tetrol: ^1^H NMR (500 MHz, CD3OD) δ 7.18 (d, J = 8.3 Hz, 1H), 6.71 (s, 1H), 6.60 (s, 1H), 6.47 (dd, J = 8.3, 2.5, Hz, 1H), 6.30 (d, J = 2.5 Hz, 1H), 3.96 (s, 1H), 3.93 (dd, J = 11.4, 1.4 Hz, 1H), 3.69 (d, J = 11.4 Hz, 1H), 3.35 (br s, 1H), 3.02 (d, J = 15.6 Hz, 1H), 2.77 (d, J = 15.6 Hz, 1H) ppm. ^13^C NMR (125 MHz, CD3OD) δ 158.0, 155.9, 145.8, 145.5, 137.6, 132.4, 131.5, 115.7, 113.1, 112.6, 110.1, 104.4, 78.3, 71.0, 51.2, 43.1 ppm.

### 2.2 Cell culture

The non-tumorigenic mammary epithelial cell line MCF10A and the breast cancer cell lines MCF7 and MDA-MB-231 (ATCC, Manassas, VA, USA) were cultured in DMEM/F12 medium (50:50, V: V; Sigma -Aldrich, St Louis, MO, USA) supplemented with 5% fetal bovine serum (FBS) and 1% antibiotics (penicillin G/streptomycin; Gibco, Waltham, MA, USA) in a humidified atmosphere containing 5% CO 2 at 37 °C.

### 2.3 Cell viability assays

The cells were seeded at 10^5^ in a 96-well plate. After adherence overnight, cells were incubated with the medium alone or with a serial dilution of brazilin, starting with the highest concentration at 40 μM/mL for 48 h. Doxorubicin and DMSO were used as controls. Then, 20 μL of MTT solution was added to each well and mixed. After 4 h, the supernatants were removed, and 100 μl of the MTT solvent (acidified isopropanol) was added to each well to dissolve the formazan crystals. Cell viability was estimated by measuring absorbance at 630 nm using a Thermo Scientific ™ Multiskan ™ FC microplate photometer. The percentage of cell viability was calculated as a function of the absorbance ratio between the cell culture treated with brazilin and the untreated control multiplied by 100, representing the cell viability (percentage of control).

### 2.4 Morphological assessment

For all untreated and untreated MDA-MB-231, MCF7, and MCF10A cells, images were captured at 48 h with an EVOS FL optical microscope using an objective 40X.

### 2.5 Cell migration assays

As described above, MDA-MB-231, MCF7, and MCF10A cells were grown to confluence in 60 mm culture plates supplemented with DMEM/F12. Cells were starved for 24 h in DMEM/F12 without FBS and treated for 2 h with Cytosine β-D-Arabinofuranoside (AraC) to inhibit cell proliferation during the experiment. After starvation, cells were scratched using a sterile 200 μL pipet tip, suspended cells were removed by washing with PBS twice, and cultures were re-fed with DMEM/F12 in the presence or absence of brazilin (0-40 mM). The progress of cell migration into the wound was monitored at 0 and 24 h using an EVOS FL microscope with a 10x objective. Cell cultures were photographed immediately after wounding and 24 h after the treatments; five images were analyzed per plate. The distance between the wound edges was measured at 0 and 24 h. The migration area was determined by measuring the total wound area using ImageJ software.

### 2.6 Molecular docking

The structure of brazilin is generated using ChemDraw (version 16.0.1.4), and prediction and screening of biological activity was performed using the “PASS Online” web resource (http://www.way2drug.com/dr/predict.php)[24]. The three-dimensional (3D) structure of the Janus kinase (JAK) 1 (4EHZ), JAK2 (4F08) and inducible nitric oxide synthase (iNOS) (1M8D) protein was obtained from the PDB database (https://www.rcsb.org), co-crystallized ligands were identified and removed from the crystall structures.

Molecular docking studies were performed with the USCF Chimera alpha (version 1.14) Autodock Vina software tool (https://www.cgl.ucsf.edu/chimera/download.html). The model was prepared by optimizing torsion angles and removal of water and ions, addition of polar hydrogen, and charge assignment. Active sites were defined using appropriately sized grid boxes around bound cocrystalline ligands. The potential binding site for the JAK1 protein, a grid was generated to identify the xyz coordinates (X = 15, Y = 4 and Z = 1), and the grid size was set to 10 × 10 × 10 Å, JAK2 with xyz coordinates (10,-52,-22), with a 15× 15 × 15 Å cube, for the iNOS protein, xyz coordinates (X = 125, Y = 112 and Z = 30), the site of coupling for the structures of the ligands was defined by establishing a cube with the dimensions 15×15×15 Å.

### 2.7 Statistical analysis

The results are presented as the mean ± SEM. Data were analyzed statistically by one-way ANOVA and Dunnett’s multiple comparison test performed comparisons. A statistical probability of P< 0.05 was considered significant.

## 3. Results and Discussion

In recent years, searching for and identifying new molecules from natural sources has been a major challenge, enabling the development of next-generation drugs to resolve global public health problems, including cancer [25]. In this regard, brazilin, a homo isoflavonoid, has acquired great importance since it can inhibit several kinases and act as an antagonist of membrane receptors involved in activating signaling pathways related to cancer [26].

### 3.1 Chemistry

Isoflavonoids have shown anti-cancer activity in several cancer cell lines, animal models, and cancer patients [19]. In our study, the ground heartwood of *Haematoxylum brasiletto* was used to obtain the compound homo isoflavonoid brazilin; 600 mg of brazilin was obtained from 1500 g of sample, with an efficiency of 0.04% (Fig. 1).

**Fig. 1.**
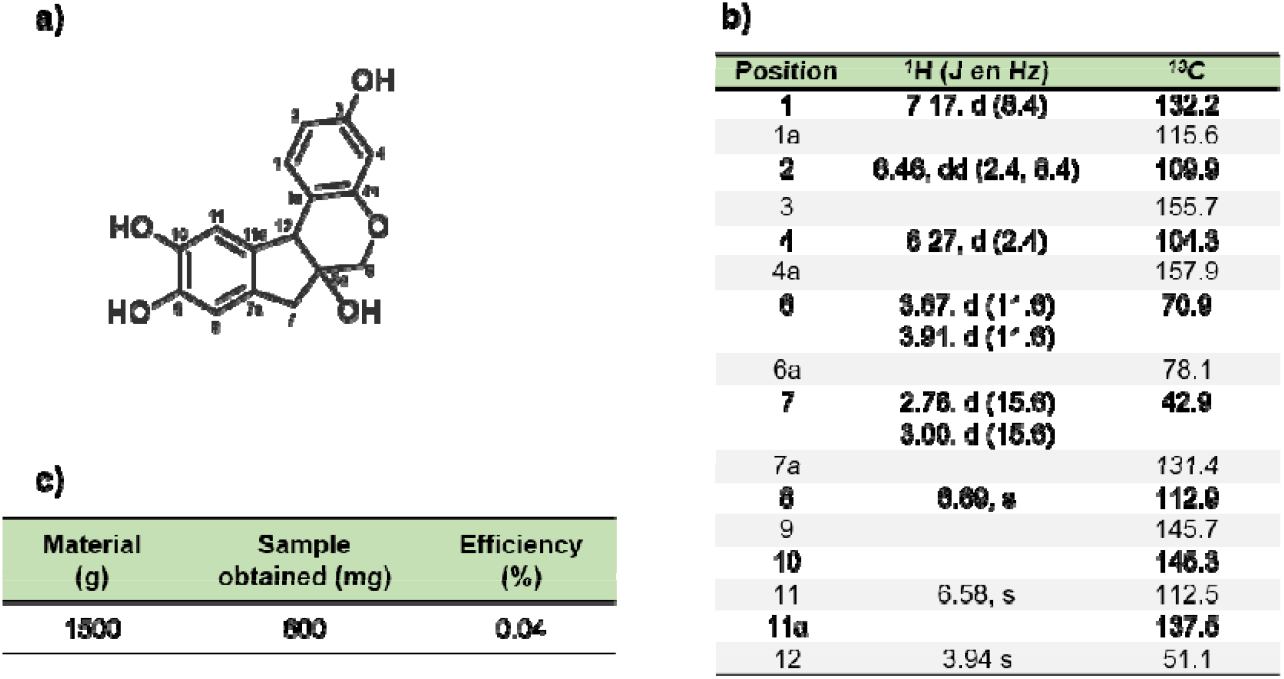
Brazilin extracted from Haematoxylum brasiletto. a) chemical structure of brazilin; b) spectroscopic da (1H RNM and 13C NMR) of brazilin; c) purification efficiency of brazilin from the heartwood of Haematoxylum brasiletto.

Recently, our working group demonstrated that brazilin, isolated from *Haematoxylum brasiletto*, decreased cell viability in different cancer cell types, including MDA-MB-231 breast cancer cells [27]. In this study, we aimed to deepen the effect of brazilin on three cell lines, the non-tumorigenic mammary epithelial cell line MCF10A, the non-invasive breast cancer cell line MCF7, and the highly invasive triple-negative cell line MDA-MB-231.

### 3.2 Morphology and viability cell assays

The evaluation of the cell morphology of a subpopulation of MDA-MB-231 cell lines changed from a mesenchymal morphology to a spherical morphology after 24 h and 48 h. Particularly, in MCF7 cells, we also observed that brazilin treatment promotes the presence of rounded cells, suggestive of dead cells (Fig. 2a), and we observed statistically significant differences of the brazilin treatments with respect to the control (untreated cells) (Fig. 2b). However, further studies are required to test this hypothesis.

**Fig. 2.**
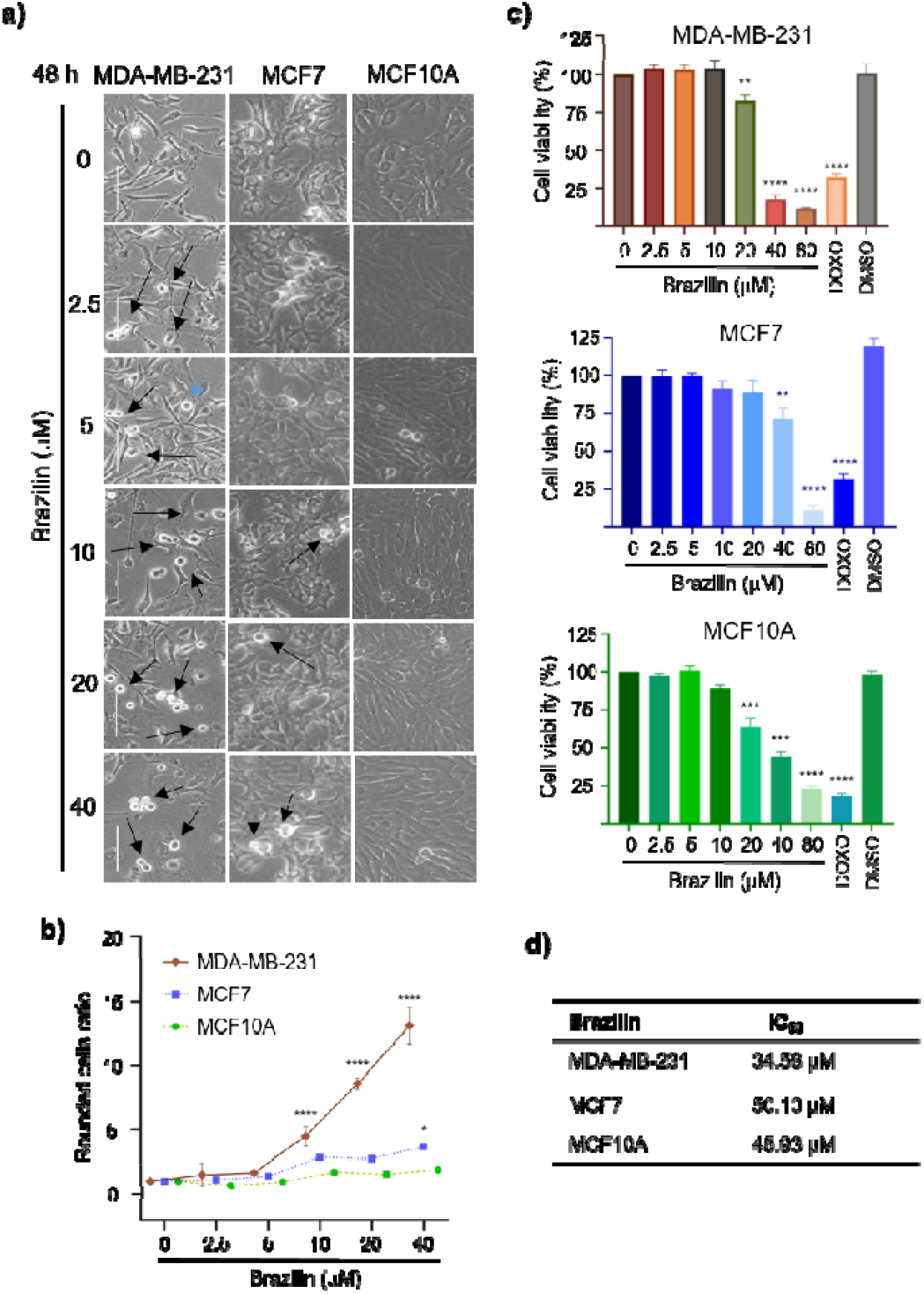
Evaluation of cell morphology and viability of breast cancer cell lines. a) Morphological appearance of breast cancer cells MDA-MB-231 and MCF7 and the non-tumorigenic breast epithelial cell line MCF10A treated with brazilin for 48 h. Arrows indicate cells with a rounded morphology after brazilin treatment. b) Cell viability of breast cancer cells and MCF10A cells after treatment with brazilin for 48 h. Differences from the control were determined with one-way ANOVA and Dunnett’s multiple comparison test. Statistical significance: *P < 0.05), **P < 0.01), ***P < 0.001), ****P < 0.0001).

Interestingly, in the non-tumor cell line MCF10A we did not observe rounded cells with brazilin treatment. On the other hand, the effect of brazilin on cell viability against the three mammary epithelium cell lines showed a selectivity in the decrease of viability in cancer cells, where a lower percentage of viability was observed in the MDA-MB-231 cell line with 47.8%, and the MCF-7 cell line 57.2% at 40 μM concentrations (Fig. 2c). We also reported that the IC50 for the three cell lines, reporting values of 34.58 μM for MDA-MB-231, 50.10 μM for MCF7 and 45.93 μM for the MCF10A cell line (Fig. 2d).

### 3.3 Cell migration

Cell migration is an essential process during the invasion and metastasis of cancer cells and is a very critical target in cancer therapy [28]. For this reason, we decided to evaluate the effect of brazilin on mammary cancer cell migration by wound closure assays (Fig. 3). Fig. 3a shows the representative photographs at 0 and 24 h in brazilin-treated MDA-MB-231 cells. In the right panel corresponds to the graph of cell migration, where brazilin 20 μM decreases cell migration by around 21%. Interestingly, brazilin 40 decreases cell migration by approximately 47% concerning the control.

**Fig. 3.**
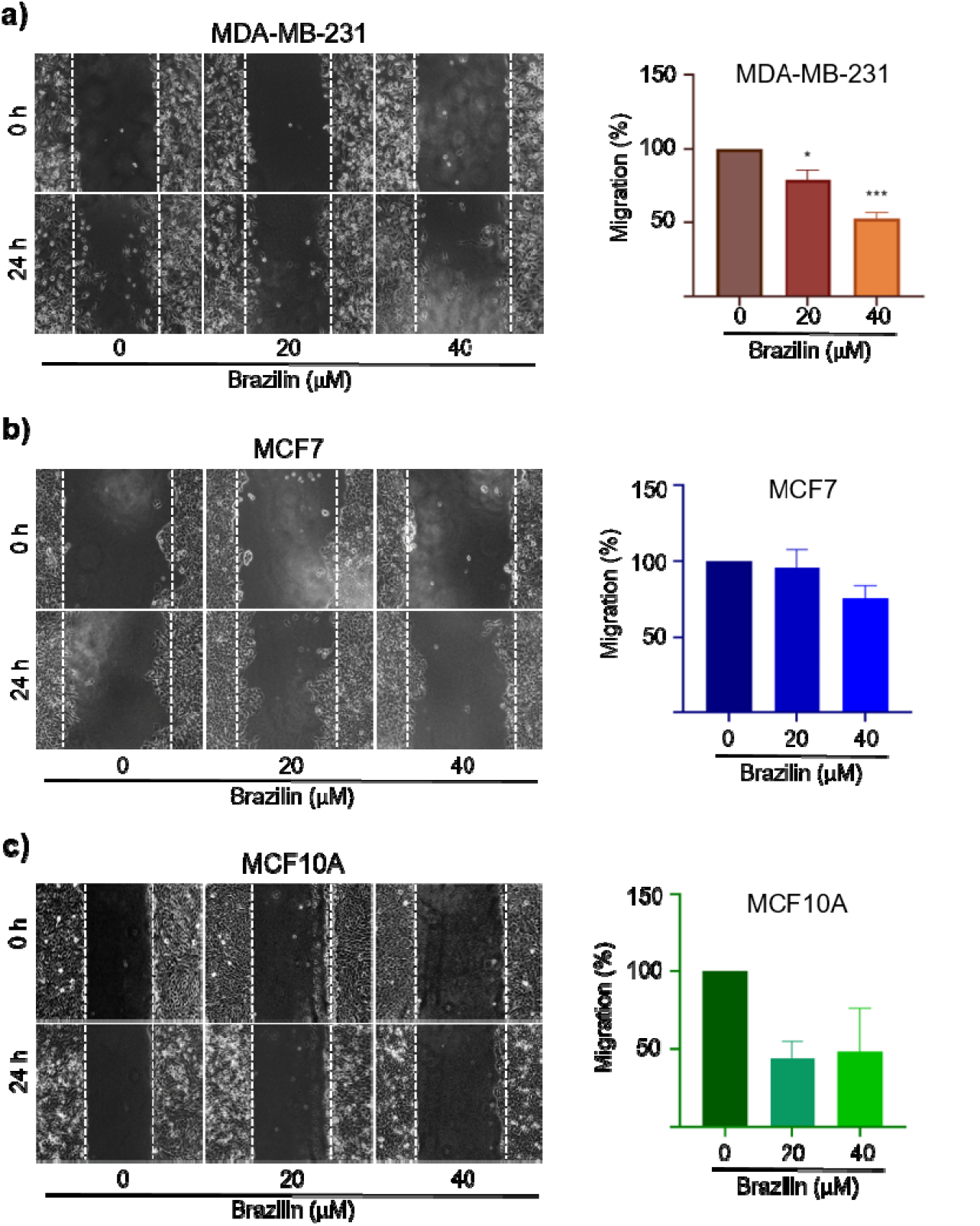
Wound closure assays in brazilin-treated breast cancer cells. a) confluent monolayers of MDA-MB-231, b) MCF7, and c) non-tumorigenic MCF10A breast cancer cells were pretreated for 2 h with Ara-C to inhibit proliferation and treated with 20 μM and at 40 μM brazilin for 24 h. The graphs represent the quantification of the percentage of cell migration for each condition. The image is representative of three independent experiments. Differences from the control were determined with one-way ANOVA and Dunnett’s multiple comparison test. Statistical significance: *P < 0.05), ***P < 0.001).

Likewise, in MCF-7 cells, brazilin 20 μM decreases cell migration, corresponding to only 4%. However, in brazilin 40 μM, cell migration decreased by 25% compared to the control (Fig. 3b). On the other hand, the photographs corresponding to MCF10A cells are shown in panel 3c, and the graph of the result analysis in right panel. Interestingly, in both concentrations with brazilin there is a decrease of about 50% in cell migration (55 and 51 %, respectively).

Recently, brazilin, isolated from *Caesalpinia sappan*, was described to inhibit cell migration, invasion, and metastasis in breast cancer [29]. On the other hand, it has been described that brazilin, in combination with doxorubicin, has a cytotoxic and anti-migratory effect on MCF7 breast cancer cells [30].

Focal adhesion kinase (FAK) is a protein involved in regulating cell migration in cancer and requires the formation of focal adhesions, a process regulated by adhesion molecules such as integrins and by various intracellular proteins [31]. FAK is a kinase that regulates the assembly and disassembly of adhesions during cell migration, and its activation is related to the autophosphorylation of the Y397; in addition, its expression is related to tumor progression and the clinical prognosis [32]. Our study found that FAK phosphorylation levels decrease by over 50% at brazilin 40 μM in MDA-MB-231 cells. However, in MCF-7 and MCF10A cells, FAK activation is not decreased (Fig. 4).

**Fig. 4.**
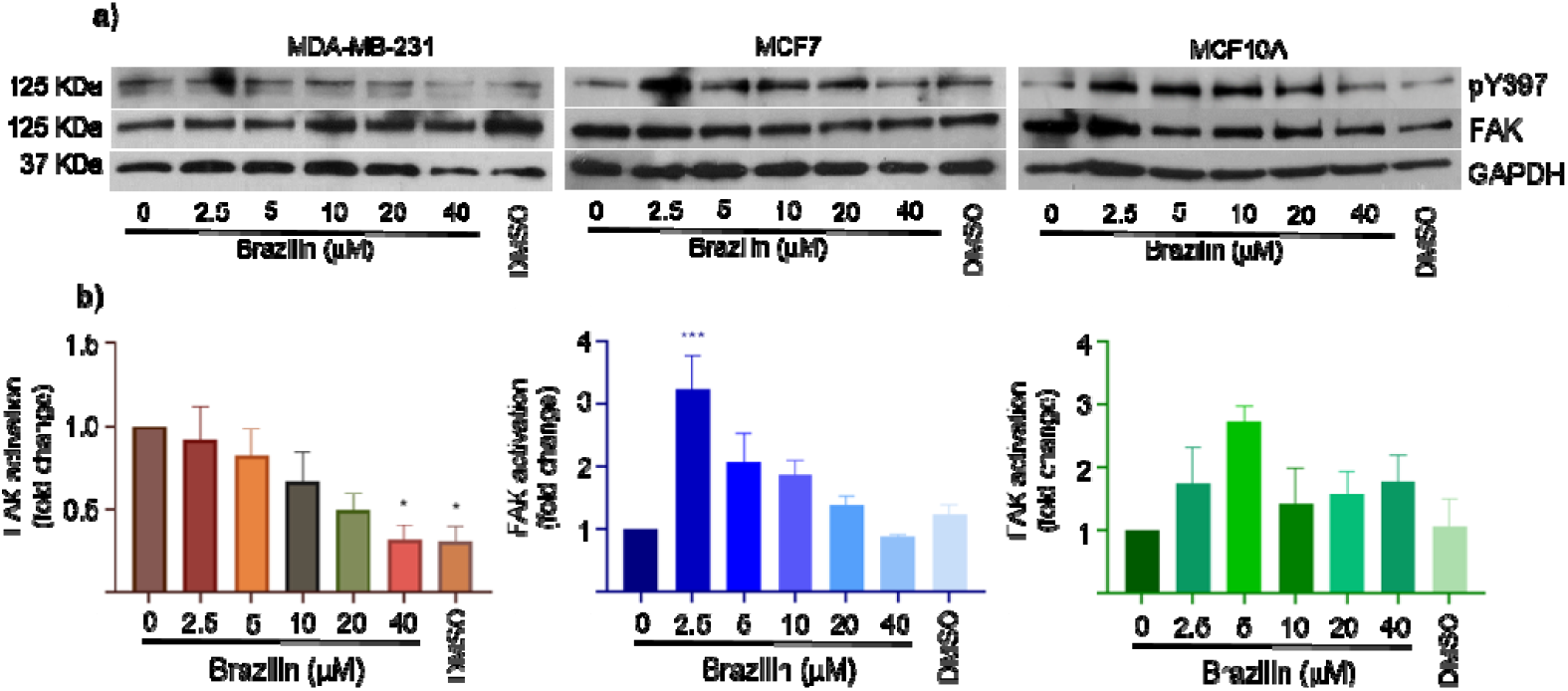
Effect of brazilin on FAK activation in MDA-MB-231, MCF-7, and MCF10A cells. Cell cultures were treated with brazilin 0, 2.5, 5, 5, 10, 20, and 40 μM and 1 μL of DMSO/mL of culture medium as control (vehicle) for 24 h. a) FAK Western blot corresponding to brazilin treatments in MDA-MB-231, MCF-7, and MCF10A cells. b) Graphs corresponding to densitometric and statistical analysis of the bands obtained by WB. Differences from the control were determined with one-way ANOVA and Dunnett’s multiple comparison test. Statistical significance: *P < 0.05), ***P < 0.001).

These findings could be explained because structurally, isoflavonoids show similarities to estrogens, bind to estrogen receptors with preferential affinity for estrogen receptor beta (ERβ), compete with 17β-estradiol for the ligand-binding domain of the receptor, and exhibit both estrogenic and anti-estrogenic activities [33,34]. In this regard, estrogen, through the ER receptor, has been reported to promote FAK activation in ovarian cancer cells [35]. However, these results should be studied in depth to establish the observed behavior.

### 3.4 Molecular docking

The biological processes observed in this study, such as cell viability or migration, maybe the result of the interaction of brazilin with Janus kinase (JAK) and iNOS proteins. Both molecules have been associated with essential proteins as signal transducers and activators of transcription, which in turn regulate the expression of various proteins involved in the proliferation, survival, migration, and invasion of different cancer cells. Interestingly, the bioinformatics data obtained here, brazilin, showed a higher probability of interaction with JAK and iNOS [26]. The iNOS protein displays the sites of interaction with Trp 366 and Cys 194, and the interaction region is characteristic of the catalytic regions (Fig. 5, Table 1)) [27], [28].

**Fig. 5.**
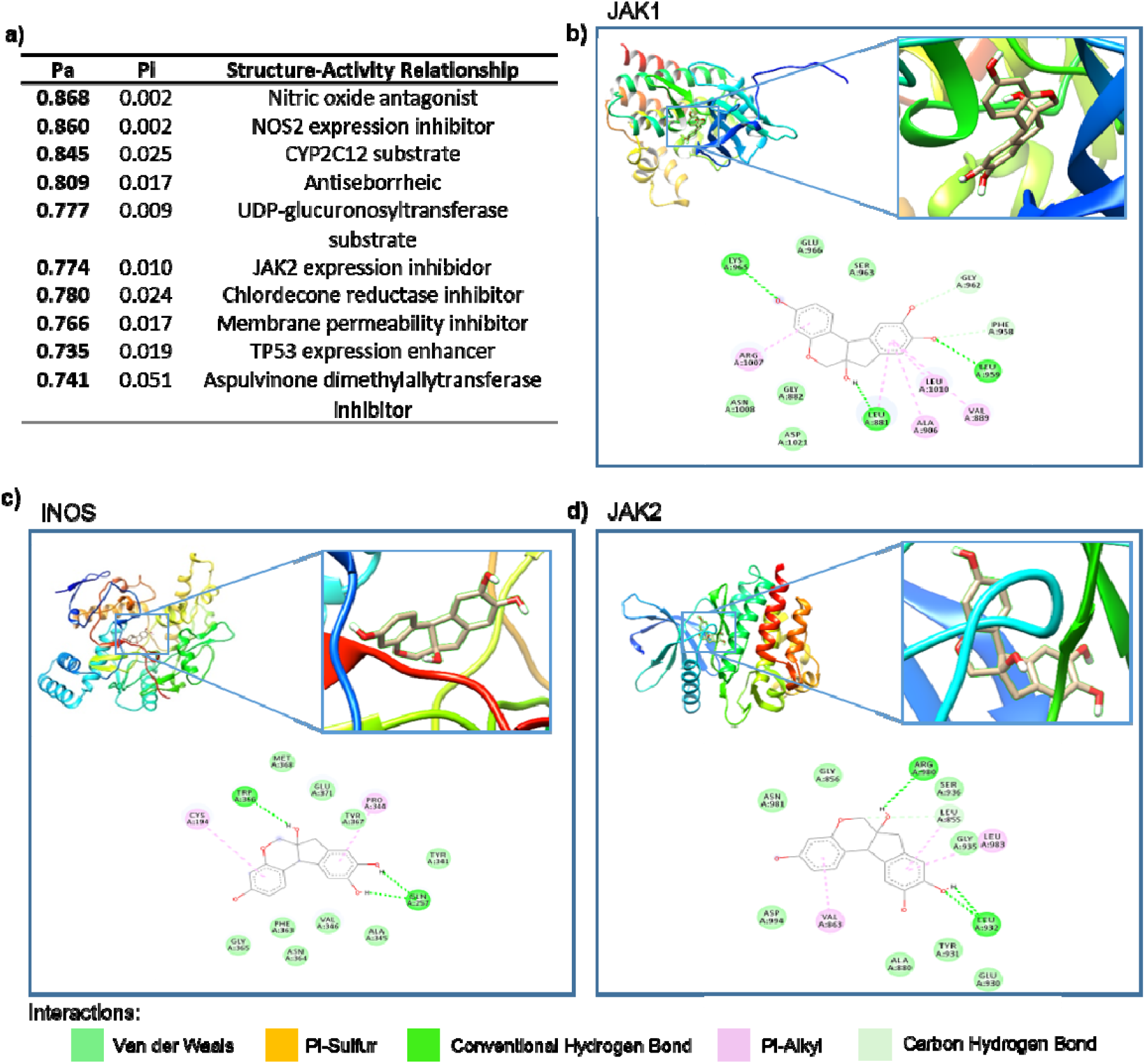
SAR and interaction analysis of brazilin. a) Description of biological targets of brazilin by structure-activity relationship prediction analysis using PASS online server. 3D protein-ligand binding models and interaction diagram by hydrogen bonding and hydrophobic interaction between brazilin and b) JAK1 (4EHZ), c) JAK2 (4F08), and d) iNOS (1M8D) proteins.

**Table 1.**
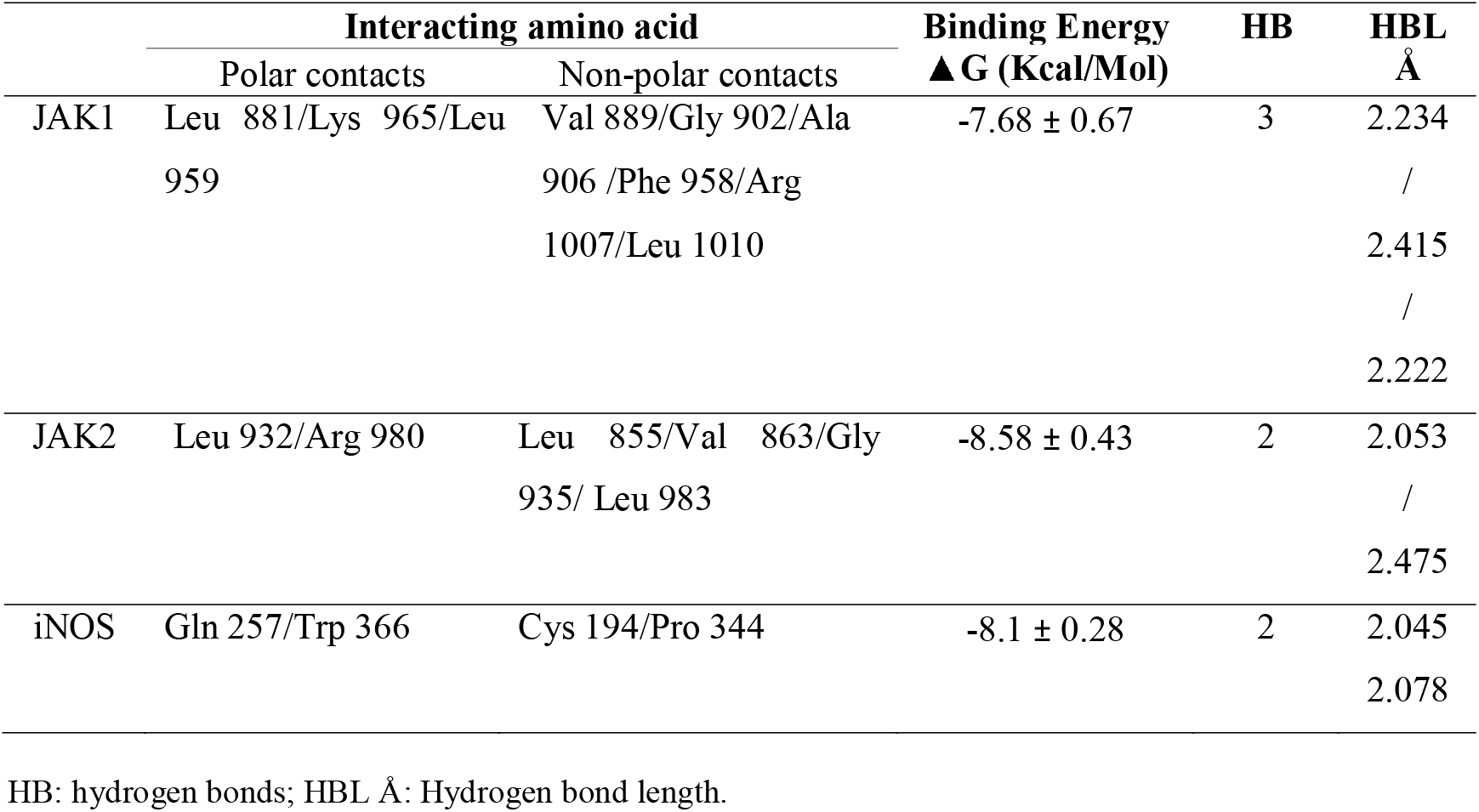
Profile of molecular docking of brazilin with JAK1, JAK2 and iNOS.

The JAK family is an essential part of the JAK/STAT signaling pathway; this pathway transduces signals from cytokines, interleukins, and growth factors acting through several transmembrane receptor families such as interleukin-6 receptor (IL-6R), leptin receptor (LEPR), epidermal growth factor receptor (EGFR), and vascular endothelial growth factor receptor (VEGFR). The intracellular domains of these receptors are associated with JAK. Ligand binding produces conformational changes in the receptors that alter the conformation of the receptor-associated JAK. Active JAK phosphorylates tyrosine residues in the cytoplasmic region of the receptor and creates binding sites that recruit signal transducer and activator of transcription (STAT). Active STAT form dimers and translocate to the nucleus when phosphorylated on tyrosine residues. The dimers bind to specific promoter sequences and modulate the transcription of genes that control cellular processes, such as proliferation, epithelial-mesenchymal transition (EMT), cell migration, differentiation, and apoptosis [29].

In uncontrolled processes due to inhibiting the transcription of genes that regulate this pathway, it is described that STAT3 is phosphorylated in 40% of breast cancer cases [30]. This phosphorylation triggers the expression of Bcl-2, Bcl-xL, Survivin, cyclin D1, c-Myc, and Mcl-1, genes promoting angiogenesis, HIF-1α, and VEGF, and genes associated with the EMT, vimentin, Twist, matrix metalloprotease (MMP)-9, and MMP-7, as well as promoting the transcription of the iNOS protein, other pathways such as NF-νB and mitogen-activated protein kinase (MAPK) contribute to the expression of the iNOS gene [30].

The iNOS enzyme allows the production of nitric oxide (NO) in cells under normal physiological conditions from the conversion of l-arginine to l-citrulline [31]. Increased NO production can promote tumor progression and metastasis by directly inducing proliferation, cell migration, and invasion of tumor cells [33,34]. Indirectly, through the expression of angiogenic factors in tumor cells, NO induces accumulation of the p53 mutation, transactivation of the EGFR by S-nitrosylation, and inhibition of integrin expression [35]. Furthermore, brazilin promotes growth inhibition and apoptosis through caspase-3 activation and subsequent PARP cleavage in human U87 glioblastoma cells and U266 myeloma cells [22,45]. On the other hand, brazilein, a product obtained by oxidation of brazilin, suppresses the expression and activity of MMP-2, which could be correlated with the inactivation of the p38 MAPK, PI3K/Akt, and NF-νB pathway [46], and as an inhibitor of P-glycoprotein (Pgp) in MCF7 cells [23], a molecule associated with multidrug resistance [47,48].

## 4. Conclusion

The observed effects of brazilin in this study could be related to flavonoid-protein interactions, which, coupled with their antioxidant properties, suggest that they may inhibit important cancer biological processes such as cell migration. Therefore, further studies, especially focused on signaling pathways and more complex models including *in vivo* models, are essential to fully define the potential of brazilin in cancer biology. Thus, brazilin could be positioned as a potential drug as a promising therapeutic option in breast cancer.

## Authors’ contributions

**Napoleon Navarro-Tito, Alberto Hernández-Moreno:** Investigation, writing the initial draft, reviewing & editing the manuscript, visualization with the figures. **Jorge Bello-Martinez, Alberto Hernández-Moreno, Napoleón-Navarro Tito**: Conceptualization, investigation. **Napoleón Navarro-Tito, Jorge Bello-Martínez:** acquisition of the financial support for the project leading to this publication. **Dania A. Nava-Tapia, Monserrat Olea-Flores, Miriam D. Zuñiga-Eulogio, Tadeo Hernández-Moreno:** Methodology, manuscript writing and revision the manuscript, editing the manuscript, visualization with the figures. Final manuscript read and approved by all authors.

## Acknowledgments

Alberto Hernandez-Moreno, Dania A. Nava Tapia, and Miriam D. Zuñiga-Eulogio were supported by the CONACYT Predoctoral Training Grant.

## Data availability

Data will be made available on request.

## References

[1] H. Sung, J. Ferlay, R.L. Siegel, M. Laversanne, I. Soerjomataram, A. Jemal, F. Bray, Global Cancer Statistics 2020: GLOBOCAN Estimates of Incidence and Mortality Worldwide for 36 Cancers in 185 Countries, CA Cancer J Clin. 71 (2021) 209–249. https://doi.org/10.3322/caac.21660.

[2] E. Laporta, J. Welsh, Modeling vitamin D actions in triple negative/basal-like breast cancer, Journal of Steroid Biochemistry and Molecular Biology. (2013) 1–9. https://doi.org/10.1016/j.jsbmb.2013.10.022.

[3] A.M. Nectoux, C. Abe, S.W. Huang, N. Ohno, J. Tabata, Y. Miyata, K. Tanaka, T. Tanaka, H. Yamamura, T. Matsui, Absorption and Metabolic Behavior of Hesperidin (Rutinosylated Hesperetin) after Single Oral Administration to Sprague-Dawley Rats, J Agric Food Chem. 67 (2019) 9812–9819. https://doi.org/10.1021/acs.jafc.9b03594.

[4] R.A.W. Christine L. Chaffer, A Perspective on Cancer Cell Metastasis, (2011) 1559–1565.

[5] R.A.W. Scott Valastyan, Tumor Metastasis=: Molecular Insights and Evolving Paradigms, 147 (2012) 275–292. https://doi.org/10.1016/j.cell.2011.09.024.Tumor.

[6] M. Kamińska, T. Ciszewski, B. Kukiełka-budny, T. Kubiatowski, Life quality of women with breast cancer after mastectomy or breast conserving therapy treated with adjuvant chemotherapy, 22 (2015) 724–730. https://doi.org/10.5604/12321966.1185784.

[7] K. Balaji, M. Sc, B. Subramanian, P. Yadav D. Ph, Radiation therapy for breast cancer=: Literature review Medical Dosimetry Radiation therapy for breast cancer=: Literature review, Medical Dosimetry. 41 (2016) 253–257. https://doi.org/10.1016/j.meddos.2016.06.005.

[8] M.S.U. Hassan, J. Ansari, D. Spooner, S.A. Hussain, Chemotherapy for breast cancer (Review), (2010) 1121–1131. https://doi.org/10.3892/or.

[9] M.I. Nounou, F. Elamrawy, N. Ahmed, K. Abdelraouf, S. Goda, H. Syed-sha-qhattal, Breast Cancer=: Conventional Diagnosis and Treatment Modalities and Recent Patents and Technologies, 9 (2015) 17–34. https://doi.org/10.4137/BCBCR.S29420.TYPE.

[10] J.B. Jia, C. Lall, T. Tirkes, R. Gulati, Chemotherapy-related complications in the kidneys and collecting system=: an imaging perspective, (2015) 479–487. https://doi.org/10.1007/s13244-015-0417-x.

[11] A. Grigorian, C.B.O. Brien, Review Article Hepatotoxicity Secondary to Chemotherapy, 2 (2014) 95–102.

[12] K.H. Schmitz, T. Disipio, L.G. Gordon, S.C. Hayes, Adverse breast cancer treatment effects=: the economic case for making rehabilitative programs standard of care, (2014). https://doi.org/10.1007/s00520-014-2539-y.

[13] A. Bauer, Industrial natural product chemistry for drug discovery and development, (2013). https://doi.org/10.1039/c3np70058e.

[14] S. Mathur, C. Hoskins, Drug development=: Lessons from nature ( Review ), (2017) 612–614. https://doi.org/10.3892/br.2017.909.

[15] J.A. Beutler, Natural Products as a Foundation for Drug Discovery THE USE OF NATURAL PRODUCTS, (2009) 1–21. https://doi.org/10.1002/0471141755.ph0911s46.

[16] D.J. Newman, G.M. Cragg, Natural Products as Sources of New Drugs over the Nearly Four Decades from 01/1981 to 09/2019, J Nat Prod. 83 (2020) 770–803. https://doi.org/10.1021/acs.jnatprod.9b01285.

[17] D.M. Kopustinskiene, V. Jakstas, A. Savickas, J. Bernatoniene, Flavonoids as anticancer agents, Nutrients. 12 (2020). https://doi.org/10.3390/nu12020457.

[18] V.M. Patil, N. Masand, Anticancer Potential of Flavonoids: Chemistry, Biological Activities, and Future Perspectives, in: Studies in Natural Products Chemistry, Elsevier B.V., 2018: pp. 401–430. https://doi.org/10.1016/B978-0-444-64179-3.00012-8.

[19] L. Cayetano-Salazar, M. Olea-Flores, M.D. Zuñiga-Eulogio, C. Weinstein-Oppenheimer, G. Fernández-Tilapa, M.A. Mendoza-Catalán, A.E. Zacapala-Gómez, J. Ortiz-Ortiz, C. Ortuño-Pineda, N. Navarro-Tito, Natural isoflavonoids in invasive cancer therapy: From bench to bedside, Phytotherapy Research. 35 (2021) 4092–4110. https://doi.org/10.1002/ptr.7072.

[20] J.L.B. Zanin, B.A. De Carvalho, P.S. Martineli, M. Henrique, J.H.G. Lago, P. Sartorelli, C. Viegas, M.G. Soares, The Genus Caesalpinia L. (Caesalpiniaceae): Phytochemical and Pharmacological Characteristics, (2012) 7887–7902. https://doi.org/10.3390/molecules17077887.

[21] R.-Z.RE. Bello-Martínez J, Jiménez-Estrada M, Rosas-Acevedo JL, Avila-Caballero LP, Vidal-Gutierrez M, Patiño-Morales C, Ortiz-Sánchez E, Antiproliferative activity of Haematoxylum brasiletto H. Karst, Pharmacogn Mag. 13 (2017) S289–S293.

[22] D. Lee, M. Lee, G. Kim, H. Noh, M. Lee, Brazilin Inhibits Growth and Induces Apoptosis in Human Glioblastoma Cells, (2013) 2449–2457. https://doi.org/10.3390/molecules18022449.

[23] N.P.L. Laksmiani, E.D.Y. Meiyanto, R.A. Susidarti, CYTOTOXIC ACTIVITY OF BRAZILEIN ISOLATED FROM SECANG (CAESALPINIA SAPPAN L .) AGAINST MCF7 / DOX CELLS BY INHIBITION OF P-GLYCOPROTEIN, 9 (2017).

[24] D.A. Filimonov, A.A. Lagunin, T.A. Gloriozova, A. V. Rudik, D.S. Druzhilovskii, P. V. Pogodin, V. V. Poroikov, Prediction of the Biological Activity Spectra of Organic Compounds Using the Pass Online Web Resource, Chem Heterocycl Compd (N Y). 50 (2014) 444–457. https://doi.org/10.1007/s10593-014-1496-1.

[25] A.G. Atanasov, S.B. Zotchev, V.M. Dirsch, I.E. Orhan, M. Banach, J.M. Rollinger, D. Barreca, W. Weckwerth, R. Bauer, E.A. Bayer, M. Majeed, A. Bishayee, V. Bochkov, G.K. Bonn, N. Braidy, F. Bucar, A. Cifuentes, G. D’Onofrio, M. Bodkin, M. Diederich, A.T. Dinkova-Kostova, T. Efferth, K. El Bairi, N. Arkells, T.P. Fan, B.L. Fiebich, M. Freissmuth, M.I. Georgiev, S. Gibbons, K.M. Godfrey, C.W. Gruber, J. Heer, L.A. Huber, E. Ibanez, A. Kijjoa, A.K. Kiss, A. Lu, F.A. Macias, M.J.S. Miller, A. Mocan, R. Müller, F. Nicoletti, G. Perry, V. Pittalà, L. Rastrelli, M. Ristow, G.L. Russo, A.S. Silva, D. Schuster, H. Sheridan, K. Skalicka-Woźniak, L. Skaltsounis, E. Sobarzo-Sánchez, D.S. Bredt, H. Stuppner, A. Sureda, N.T. Tzvetkov, R.A. Vacca, B.B. Aggarwal, M. Battino, F. Giampieri, M. Wink, J.L. Wolfender, J. Xiao, A.W.K. Yeung, G. Lizard, M.A. Popp, M. Heinrich, I. Berindan-Neagoe, M. Stadler, M. Daglia, R. Verpoorte, C.T. Supuran, Natural products in drug discovery: advances and opportunities, Nat Rev Drug Discov. 20 (2021) 200–216. https://doi.org/10.1038/s41573-020-00114-z.

[26] D.A. Nava-Tapia, L. Cayetano-Salazar, L.D. Herrera-Zúñiga, J. Bello-Martínez, M.A. Mendoza-Catalán, N. Navarro-Tito, Brazilin: Biological activities and therapeutic potential in chronic degenerative diseases and cancer, Pharmacol Res. 175 (2022). https://doi.org/10.1016/j.phrs.2021.106023.

[27] J. Bello-Martínez, M. Jiménez-Estrada, J.L. Rosas-Acevedo, L.P. Avila-Caballero, M. Vidal-Gutierrez, C. Patiño-Morales, E. Ortiz-Sánchez, R.E. Robles-Zepeda, Antiproliferative activity of Haematoxylum brasiletto H. Karst, Pharmacogn Mag. 13 (2017) S289–S293. https://doi.org/10.4103/pm.pm_466_16.

[28] C.H. Stuelten, C.A. Parent, D.J. Montell, Cell motility in cancer invasion and metastasis: Insights from simple model organisms, Nat Rev Cancer. 18 (2018) 296–312. https://doi.org/10.1038/nrc.2018.15.

[29] Yang X, Liang Y, Zhao L, Chen L, Yang Y, Wang J, Yan L, Zhang S, Liu X, Zhang H, Brazilin inhibits cell migration and metastasis in breast cancer, Biol. Pharm. Bull. 46 (2023) 773–780.

[30] R.I. Jenie, S. Handayani, R.A. Susidarti, L.Z. Udin, E. Meiyanto, The cytotoxic and antimigratory activity of Brazilin-doxorubicin on MCF-7/HER2 cells, Adv Pharm Bull. 8 (2018) 507–516. https://doi.org/10.15171/apb.2018.059.

[31] Maziveyi Mazvita, Alahari Suresh K., Cell matrix adhesions in cancer: The proteins that form the glue, 2017.

[32] K. Katoh, FAK-Dependent Cell Motility and Cell Elongation, Cells. 9 (2020). https://doi.org/10.3390/cells9010192.

[33] P. Basu, C. Maier, Phytoestrogens and breast cancer: In vitro anticancer activities of isoflavones, lignans, coumestans, stilbenes and their analogs and derivatives, Biomedicine & Pharmacotherapy. 107 (2018) 1648–1666. https://doi.org/10.1016/j.biopha.2018.08.100.

[34] S.R. Lepri, R.C. Luiz, L.C. Zanelatto, P.B.G. da Silva, D. Sartori, L.R. Ribeiro, M.S. Mantovani, Chemoprotective activity of the isoflavones, genistein and daidzein on mutagenicity induced by direct and indirect mutagens in cultured HTC cells, Cytotechnology. 65 (2013) 213–222. https://doi.org/10.1007/s10616-012-9476-8.

[35] Y.C. Hung, W.C. Chang, L.M. Chen, Y.Y. Chang, L.Y. Wu, W.M. Chung, T.Y. Lin, L.C. Chen, W.L. Ma, Non-genomic estrogen/estrogen receptor α promotes cellular malignancy of immature ovarian teratoma in vitro, J Cell Physiol. 229 (2014) 752–761. https://doi.org/10.1002/jcp.24495.

[36] E. Bousoik, H.M. Aliabadi, “Do We Know Jack” About JAK=? A Closer Look at JAK / STAT Signaling Pathway, 8 (2018) 1–20. https://doi.org/10.3389/fonc.2018.00287.

[37] R.J. Rosenfeld, E.D. Garcin, K. Panda, G. Andersson, A. Åberg, A. V Wallace, G.M. Morris, A.J. Olson, D.J. Stuehr, J.A. Tainer, E.D. Getzoff, Conformational Changes in Nitric Oxide Synthases Induced by Chlorzoxazone and Nitroindazoles=: Crystallographic and Computational Analyses of Inhibitor Potency †, (2002) 13915–13925.

[38] M. Zak, R. Mendonca, M. Balazs, K. Barrett, P. Bergeron, W.S. Blair, C. Chang, G. Deshmukh, J. Devoss, P.S. Dragovich, C. Eigenbrot, N. Ghilardi, P. Gibbons, S. Gradl, C. Hamman, E.J. Hanan, E. Harstad, P.R. Hewitt, C.A. Hurley, T. Jin, A. Johnson, T. Johnson, J.R. Kenny, M.F.T. Koehler, P.B. Kohli, J.J. Kulagowski, S. Labadie, J. Liao, M. Liimatta, Z. Lin, P.J. Lupardus, R.J. Maxey, J.M. Murray, R. Pulk, M. Rodriguez, S. Savage, S. Shia, M. Ste, S. Ubhayakar, M. Ultsch, A. Van Abbema, S.I. Ward, L. Xiao, Y. Xiao, Discovery and Optimization of C -2 Methyl Imidazopyrrolopyridines as Potent and Orally Bioavailable JAK1 Inhibitors with Selectivity over JAK2, (2012).

[39] J. Pencik, H. Thi, T. Pham, J. Schmoellerl, T. Javaheri, Europe PMC Funders Group JAK-STAT signaling in cancer=: From cytokines to non-coding genome, (2018) 26–36. https://doi.org/10.1016/j.cyto.2016.06.017.JAK-STAT.

[40] B. Groner, V. Von Manstein, Jak Stat signaling and cancer=: Opportunities, benefits and side effects of targeted inhibition Molecular and Cellular Endocrinology Jak Stat signaling and cancer=: Opportunities, bene fi ts and side effects of targeted inhibition, Mol Cell Endocrinol. 451 (2017) 1–14. https://doi.org/10.1016/j.mce.2017.05.033.

[41] F. Vannini, K. Kash, N. Nath, Redox Biology The dual role of iNOS in cancer $, 6 (2015) 334–343. https://doi.org/10.1016/j.redox.2015.08.009.

[42] S.A. Glynn, R.M. Stephens, S. Ambs, S.A. Glynn, B.J. Boersma, T.H. Dorsey, M. Yi, H.G. Yfantis, L.A. Ridnour, D.N. Martin, C.H. Switzer, R.S. Hudson, D.A. Wink, D.H. Lee, R.M. Stephens, S. Ambs, negative breast cancer patients Increased NOS2 predicts poor survival in estrogen receptor – negative breast cancer patients, 120 (2010) 3843–3854. https://doi.org/10.1172/JCI42059.tion.

[43] S. Granados-principal, Y. Liu, M.L. Guevara, E. Blanco, D.S. Choi, W. Qian, T. Patel, A.A. Rodriguez, J. Cusimano, H.L. Weiss, H. Zhao, M.D. Landis, B. Dave, S.S. Gross, J.C. Chang, Inhibition of iNOS as a novel effective targeted therapy against triple-negative breast cancer, (2015) 1–16. https://doi.org/10.1186/s13058-015-0527-x.

[44] E.M. Walsh, M.M. Keane, D.A. Wink, G. Callagy, A. Sharon, I. Program, Review of Triple Negative Breast Cancer and the Impact of Inducible Nitric Oxide Synthase on Tumor Biology and Patient Outcomes, 21 (2019) 333–351. https://doi.org/10.1615/CritRevOncog.2017021307.Review.

[45] B. Kim, S. Kim, S. Jeong, E.J. Sohn, J.H. Jung, M.H. Lee, S. Kim, Brazilin Induces Apoptosis and G2/M Arrest via Inactivation of Histone Deacetylase in Multiple Myeloma U266 Cells, (2012) 8–15.

[46] C.Y. Hsieh, P.C. Tsai, C.L. Chu, F.R. Chang, L. Sen Chang, Y.C. Wu, S.R. Lin, Brazilein suppresses migration and invasion of MDA-MB-231 breast cancer cells, Chem Biol Interact. 204 (2013) 105–115. https://doi.org/10.1016/j.cbi.2013.05.005.

[47] H.M. Abdallah, A.M. Al-abd, A.M. El-halawany, P-glycoprotein inhibitors of natural origin as potential tumor chemo-sensitizers=: A review, J Adv Res. 6 (2015) 45–62. https://doi.org/10.1016/j.jare.2014.11.008.

[48] O. Weso, J. Wi, Ś. Kamila, A. Krawczenko, 8-Prenylnaringenin is an inhibitor of multidrug resistance-associated transporters, P-glycoprotein and MRP1 8-Prenylnaringenin is an inhibitor of multidrug resistance-associated transporters, Pglycoprotein and MRP1, (2018). https://doi.org/10.1016/j.ejphar.2010.06.069.

